# Prevention of enteric bacterial infections and modulation of gut microbiota composition with conjugated linoleic acids producing *Lactobacillus* in mice

**DOI:** 10.1101/571117

**Authors:** Mengfei Peng, Zajeba Tabashsum, Puja Patel, Cassandra Bernhardt, Chitrine Biswas, Jianghong Meng, Debabrata Biswas

## Abstract

Probiotics are recognized to outcompete pathogenic bacteria by receptor-mediated colonizing and secreting functional metabolites which have direct antimicrobial activities towards pathogens and/or improving host’s gut health and immunity. We have constructed a *Lactobacillus casei* (LC) probiotic strain, LC^+*mcra*^, by inserting *mcra* (myosin cross-reactive antigen) gene, which stimulates the conversion of conjugated linoleic acids. In this study, we evaluated the protective roles of LC^+*mcra*^ against pathogenic *Salmonella enterica* serovar Typhimurium (ST) and enterohaemorrhagic *E. coli* (EHEC) infection in BALB/cJ mice. Through a series of *in vivo* investigation, we observed that LC^+*mcra*^ colonized efficiently in mice gut and competitively reduced the infection with ST and EHEC in various locations of small and large intestine, specifically cecum, jejunum, and ileum (*p*<0.05). The cecal microbiota in ST-challenged mice with LC^+*mcra*^ protection were positively modulated with higher relative abundances Firmicutes but lower Proteobacteria plus increased bacterial species diversity/richness based on 16S metagenomic sequencing. Based on cytokine gene expression analysis by qRT-PCR, mice pretreated with LC^+*mcra*^ were found with attenuated bacterial pathogen-induced gut inflammation. Furthermore, mice fed LC^+*mcra*^ daily for one week could protect themselves from the impairments caused by enteric infections with ST or EHEC. These impairments include weight loss, negative hematological changes, intestinal histological alterations, and potential death. This *in vivo* study suggests that daily consumption of novel conjugated linoleic acids over-producing probiotic might be efficient in improving gut intestinal microbiome composition and preventing/combating foodborne enteric bacterial infections with pathogenic *Salmonella* and diarrheagenic *E. coli.*

**Author summary:** Numerous bacteria colonize throughout the gastrointestinal tract and form a complex microbial ecosystem known as gut microbiota. A balanced microbial composition is crucial for maintaining proper gut health and host defense against pathogenic microbes. However, enteric bacterial infections could cause illness and even lead to death of host when foodborne pathogens like *Salmonella* and enterohaemorrhagic *E. coli* (EHEC) invade gut intestine and cause imbalance of gut microbiota. Beneficial microbes in gastrointestinal tract such as *Lactobacillus* and their secreted bio-active metabolites, are potential bio-agents to improve gut immunity and outcompete bacterial pathogens. In this study, to evaluate roles of novel *Lactobacillus* strain LC^+*mcra*^ which produce higher amount of a group of beneficial secondary metabolites called conjugated linoleic acids, we have shown that daily oral administration of this LC^+*mcra*^ for one-week in mice lead to higher proportion of beneficial bacterial colonization in different locations of intestine and a significant reduction of pathogenic *Salmonella* and EHEC colonization. Furthermore, mice fed with LC^*+mcra*^ restore and modulate *Salmonella* infection-induced negative impact on gut microbiota composition and protect themselves from various levels of physiological damage.

## Introduction

The majority of human gut epithelial surfaces are colonized and safeguarded by a tremendous number of microorganisms including bacteria, viruses, fungi and protozoans which are known as common gut microflora; each of them is crucial in forming and balancing a complex ecosystem with microbial diversity [1]. These large number of microorganisms build up a microbial genetic repertoire approximately 100 times greater than that of the human host. Diversity of these microbes, specifically number of diverse bacterial species, is essential for good health and immunity of host [2]. According to recent reports, human distal gastrointestinal (GI) tract can house more than 1000 distinct bacterial species, and the total number was estimated to be larger than 10^14^ CFU/gm of fecal material [3]. Bacteroidetes, Firmicutes, Proteobacteria, and Actinobacteria are the prevalent bacterial phyla in human gut microbiota and each of these phyla contains dozens of bacterial genus and hundreds of species [4–6].

In a homeostasis gut ecosystem, most of the commensal bacteria colonize and survive symbiotically, whereas conditions such as immunodeficiency, malnutrition, and antibiotic-therapy cause dysbiosis and imbalance of commensal bacteria that induce pathogenesis and cause diseases [7,8]. Furthermore, broad-spectrum antibiotic therapy or any other detrimental conditions may disturb the gut ecosystem balance long-term or lead to chronically irritated bowels, reducing the number of beneficial bacteria and increasing the number of opportunistic pathogens and their toxic products that further weaken the host defense and/or induce inflammation and damage [9]. As a consequence of imbalanced gut microflora, opportunistic pathogens, their produced metabolites, proteins, and/or toxins can take over the gut ecosystem and negatively impact host gut health.

*Salmonella* and diarrheagenic *Escherichia coli* generally infect human gut intestine through consumption of contaminated foods and/or drinks [10–12]. Once these Gram-negative enteric pathogenic bacteria arrive in host gut, their complex type III secretion systems are activated, enabling them to introduce effector proteins directly into cell cytoplasm. Series of these cascades induce systematic infections causing acute or chronic inflammation and other serious disorders in the host [13]. However, such enteric illness is usually facilitated by compromised gut immunity and dysbiotic gut microbiota which provide those enteric bacterial pathogens with weakened colonization resistance [14]. On the other hand, traditional antibiotic therapy has been found to lyse enterohemorrhagic *E. coli* (EHEC) which further increases the risk for post infectious sequelae Hemolytic-uremic syndrome (HUS) in the patients [15,16]. In such situations, procommensal strategies by application of probiotics, prebiotics, and synbiotics can be considered as priority in prevention and treatment of foodborne such bacterial pathogen-induced enteric illness [11,17,18]. With a promising scheme, it allows an establishment or recovery of the healthy enteric microbial ecosystem by introducing native, exogenous, or genetically engineered beneficial probiotics without inducing deleterious effects (like antibiotics) on human commensal gut bacteria [14,19].

Recently, we constructed and reported the role of a multi-functional *Lactobacillus casei* probiotic strain overexpressing myosin cross-reactive antigen gene (*mcra*), named as LC^+*mcra*^ [20]. Several groups of researchers have demonstrated the health-beneficial effects of conjugated linoleic acids, such as anti-carcinogenesis, anti-oxidant, and anti-microbial effects [14,21,22]. Similarly, we have also revealed the anti-pathogenic and anti-inflammatory properties of linoleic acids over-producing *L. casei* (LC^+*mcra*^) based on *in vitro* examination. Here in this study, we aimed to evaluate the protective roles of LC^+*mcra*^ on modulating/recovering gut intestinal microflora composition and combating/alleviating foodborne enteric bacterial pathogenic infections *in vivo* based on mice model.

## Results

### Probiotics preventing ST infection induced physiological abnormalities in mice

The weight of each mice was monitored every day for the purpose of investigating if probiotics preventive administration could rescue mice from weight loss due to ST/EHEC infection (Fig 1). Within the entire 4-week rearing, a total of 12 mice in control group (no probiotic given), 7 mice in group given wild-type probiotic LC strain, and 1 mouse in group given linoleic acid over-expressed mutant LC^+*mcra*^ strain were sacrificed due to their health abnormality induced by ST infection. These sacrificed individuals included 8 mice from control and 5 mice from LC treatment found self-death due to ST challenge, but none from LC^+*mcra*^ treatment, which provided us the ST survival rates as 60% in control group, 75% in LC group, and 100% in LC^+*mcra*^ group. The death of the mice was generally accompanied with extreme (>20%) weight loss to approximately 8-10 g.

**Fig 1.**
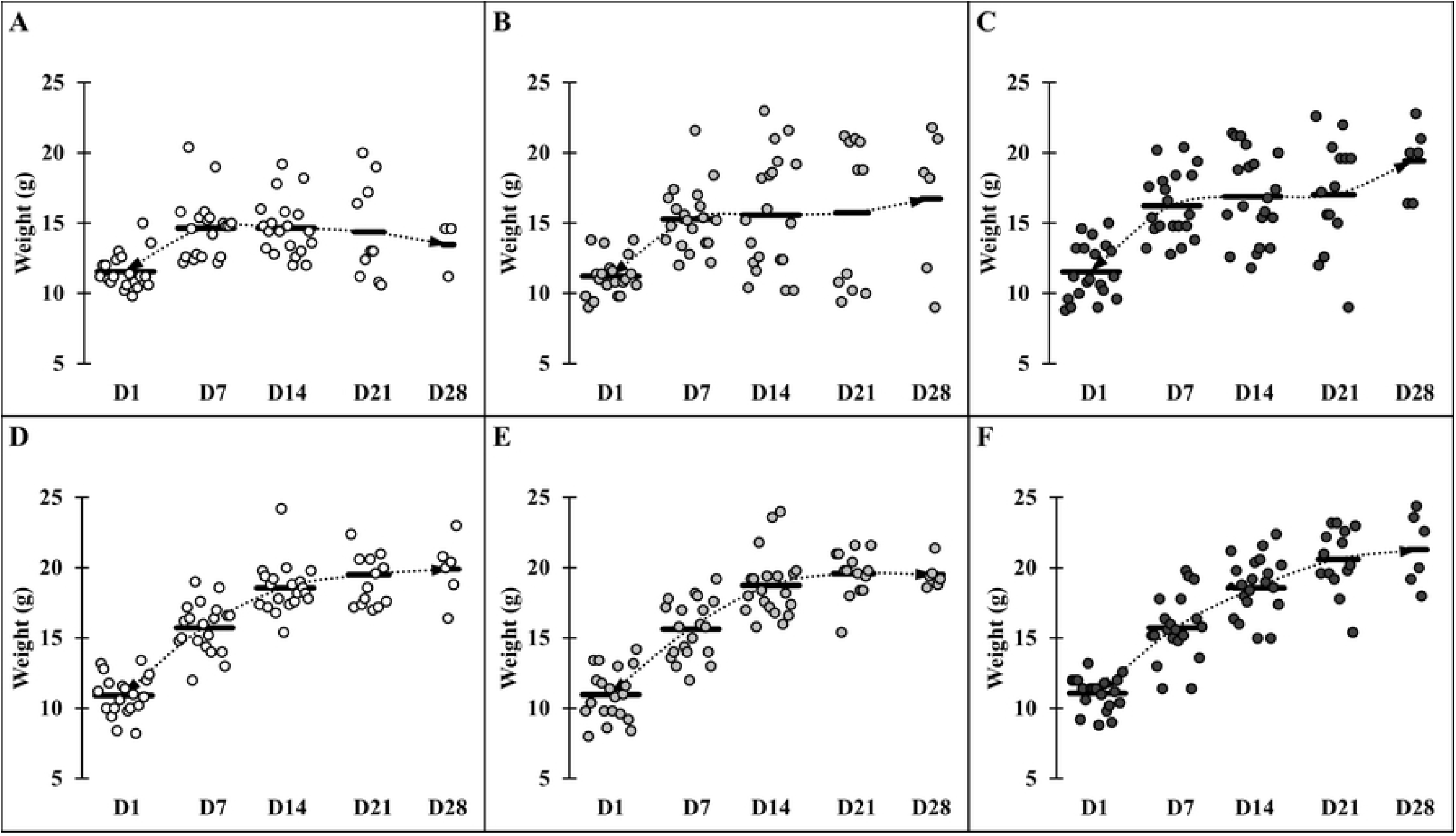
Comparative weight gain and loss in mice across different groups. Mice groups were assigned with the following manner: (A) ST infection, (B) ST infection and LC 1-week pre-treatment, (C) ST infection and LC^+*mcra*^ 1-week pre-treatment, (D) EHEC infection, (E) EHEC infection and LC 1-week pre-treatment, and (F) EHEC infection and LC^+*mcra*^ 1-week pre-treatment. Each dot indicates individual mouse weight and horizontal bars at each time point indicate averaged weight of mice in accordant group.

At the end of week 2, the average weight of mice in control group reached approximately 14-16g, whereas both groups of mice which were given either LC or LC^+*mcra*^ gained weight at range of 1-2g more compared to the control group of mice. Once mice were challenged with ST, the average weight gain trend of mice in control group which was not given probiotic was suspended and remained at 14.65 g during 1^st^ week of post-challenge. Then the weight of those mice decreased to 14.36 g and 13.47 g at 2^nd^ and 3^rd^ post-infection weeks, respectively. However, the mice which were administrated LC^+*mcra*^ kept continuing to gain average weight. In spite of the negative effect induced by ST infection, mice which were given LC^+*mcra*^ gained weight at 16.88 g, 17.02 g, and 19.12 g at the 1^st^, 2^nd^, and 3^rd^ week of post-infections, respectively. The wild-type probiotic, LC fed mice exhibited mild effects in maintaining the average body weight during the first two weeks of ST infection and gained approximately 1.5 g weight at the end of 3^rd^ post-infection week. On the other hand, we failed to observe any negative effects including average weight loss induced by EHEC infection. However, the oral administration of LC^+*mcra*^ was more effective than LC in promoting the weight earning of mice by 1.1g and 1.4 g averagely at the 2^nd^ and 3^rd^ post-infection weeks compared with control group.

### Reduction on colonization of ST and EHEC in probiotics fed mice

Either LC or LC^+*mcra*^ was orally administrated to mice in order to examine their colonization ability in mice gut and evaluate their preventive role in altering enteric pathogenic bacterial colonization and infection in gastrointestinal tract of mice using BALB/cJ mice model. According to the colonization data collected from two individual mice trials, both LC and LC^+*mcra*^ were able to colonize well in gut of BALB/cJ mice but the genetically modified probiotic strain, LC^+*mcra*^ could colonize in the mice gut more aggressively compare to the wild-type LC strain. Further, both LC and LC^+*mcra*^ significantly reduced the colonization and infection of both enteric bacterial pathogens, ST and EHEC in BALB/cJ mice. We found that mice fed with LC^+*mcra*^ could defend ST infection remarkably and recover fully within a week of challenge. Specifically, mice highly colonized with LC^+*mcra*^ strain were able to reduce significantly (approximately 1 log CFU/g) cecal colonization with ST compare to the group of mice which were given wild-type LC strain at all three time points (14, 21, and 28 d) (Fig 2A).

**Fig 2.**
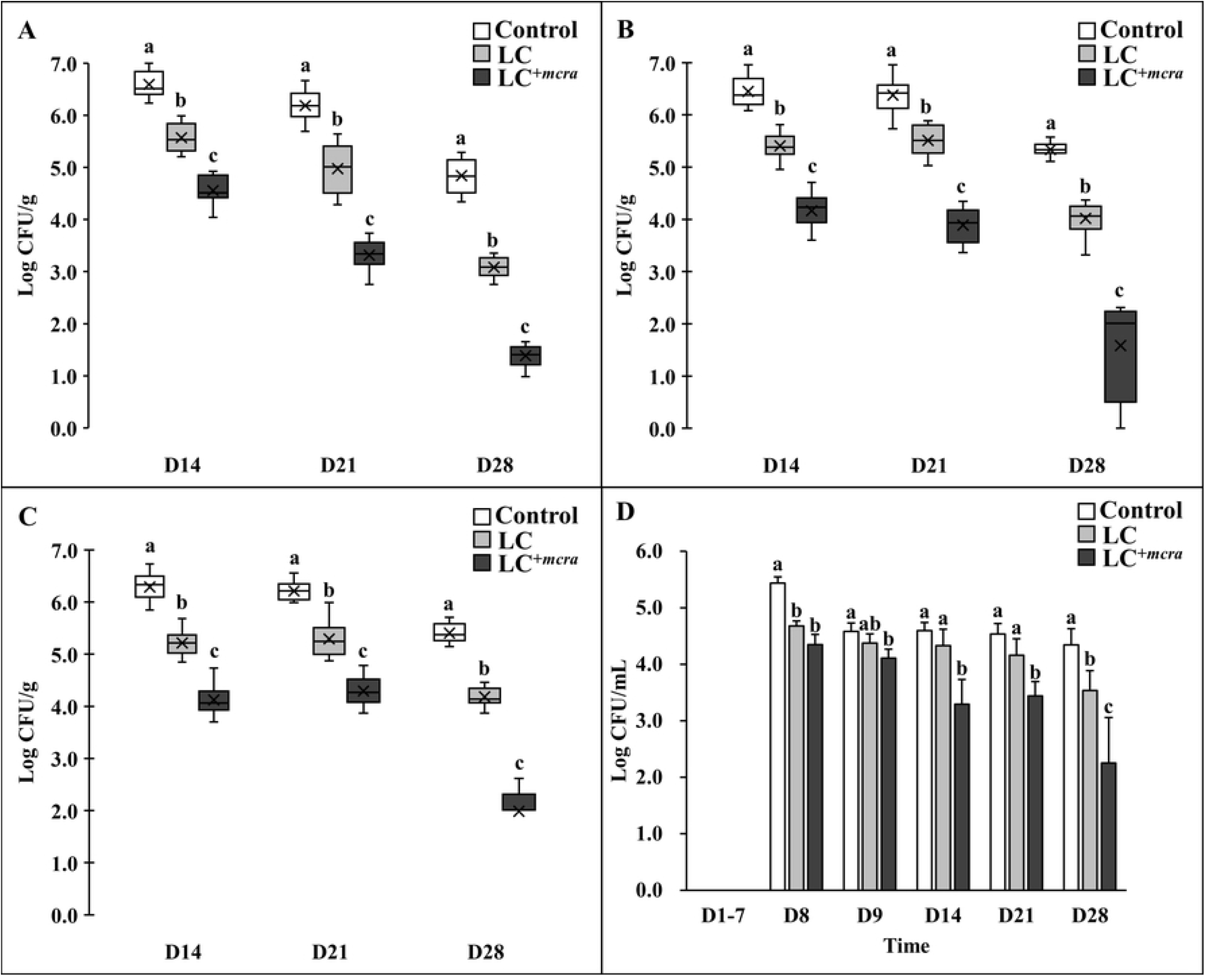
Effect of LC^+*mcra*^ on reducing colonization of ST in mice gut intestine. The bacterial numbers of ST at 14, 21, and 28 days in ileum (A), jejunum (B), cecum (C), and feces (D) from ST-infected mice with no probiotic treatment, LC, or LC^+*mcra*^ 1-week pre-treatment were investigated in triplicate. Different letters (‘a’ through ‘c’) at single time point are significantly different (p < 0.05) in the numbers of ST among control and treatments.

To compare the colonization of ST in jejunum, we observed that LC or LC^+*mcra*^ pre-administrated mice were colonized with lower number of ST at rang of 1.0 to 2.3 log CFU ST per gram jejunum fluids at 1^st^ week post-infection, 0.9 and 2.5 log CFU/g at 2^nd^ week post-infection, and 1.3 and 3.7 log CFU/g at 3^rd^ week post-infection (Fig 2B). Similarly, LC and LC^+*mcra*^ pre-administrated mice were colonized with ST in lower rate of 1.7 and 2.2 log CFU per gram ileum fluids at 1^st^ week post-infection, 0.9 and 1.9 log CFU/g on 2^nd^ week post-infection, and 1.2 and 3.4 log CFU/g on 3^rd^ week post-infection to the control mice (Fig 2C).

The significant reduction on ST gut intestinal colonization was also observed in form of decreased ST fecal shedding. On the 8^th^ day after mice were challenged with ST, both groups of mice administrated with either wild-type probiotic LC or genetically modified probiotic LC^+*mcra*^ strain were colonized with reduced number (0.8 to 1.1 log CFU/mL) ST in feces but the differences became unsubstantial at the 9^th^ day. However, notably major effectiveness of LC^+*mcra*^ started to appear in mice after 1^st^ week post-infection, at which 1.3 log CFU/mL less ST was recovered from mice feces. In the subsequent two weeks, LC^+*mcra*^ fed mice were observed with 1.1 and 2.1 log CFU/mL continuous ST reduction on fecal shedding.

On the other hand, mice which were pretreated with LC barely reduced the EHEC colonization in jejunum and ilium, whereas mice pretreated with LC^+*mcra*^ showed significant influence in EHEC colonization resistance (Fig 3). Specifically, LC^+*mcra*^ fed mice were capable of significantly reducing the colonization of EHEC at 2.3, 1.6, and 0.9 log CFU/g in cecum, 1.6, 1.8, and 2.7 log CFU/g in jejunum, and 2.8, 1.8 and 2.1 log CFU/g in ileum at the 1^st^, 2^nd^, and 3^rd^ week post-challenge. Meanwhile, consequential decreased EHEC fecal shedding was detected in LC^+*mcra*^ fed mice as well. However, only insignificant reductions (0.1 to 0.5 CFU EHEC less per mL feces) were found during the first two days after EHEC challenge on EHEC-free mice (the 8^th^ and 9^th^ day). The LC^+*mcra*^ administration substantially lowered 0.9, 1.9, and 2.2 CFU/mL EHEC fecal shedding at the 1^st^, 2^nd^, and 3^rd^ post-infection weeks in comparison with control.

**Fig 3.**
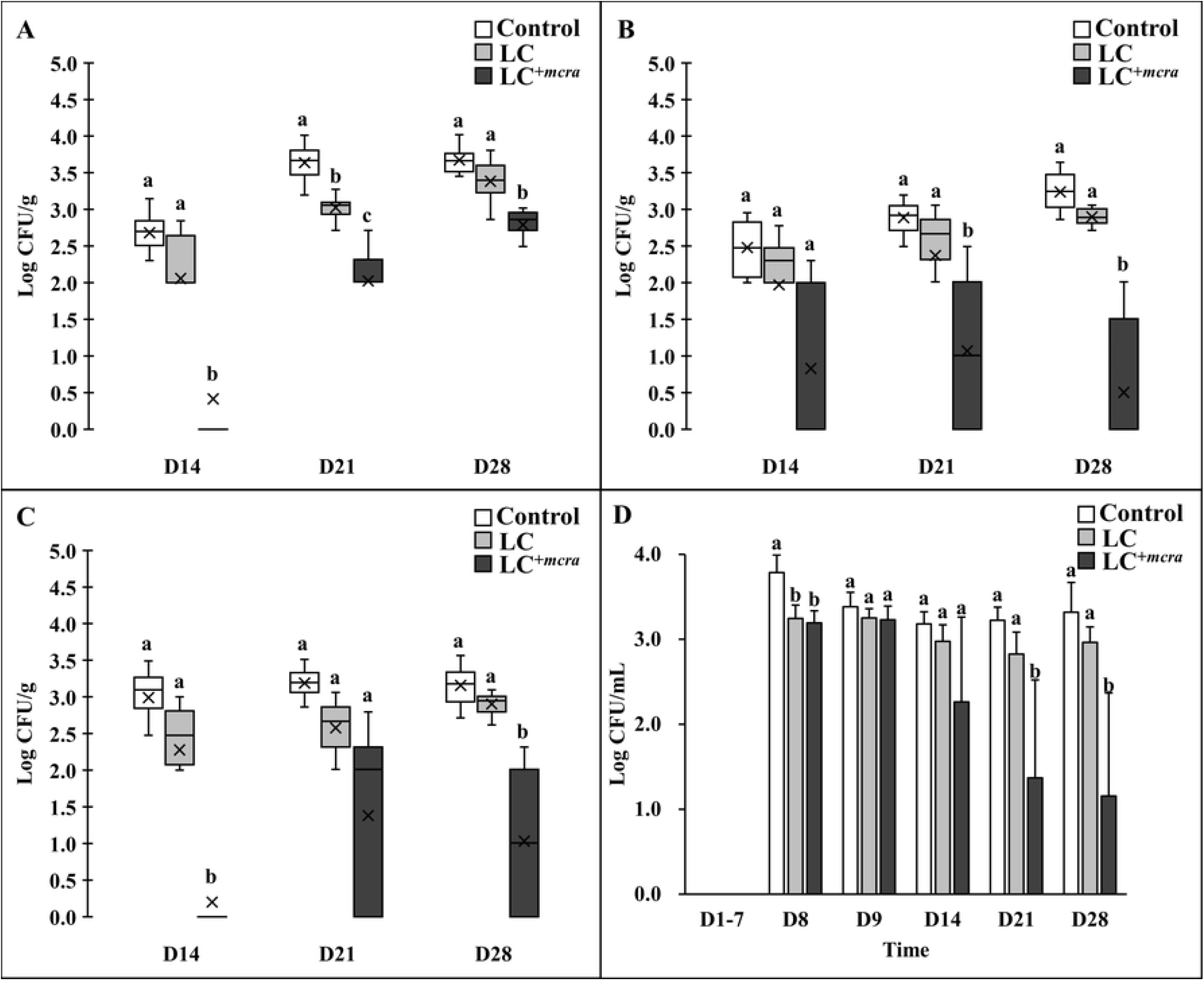
Effect of LC^+*mcra*^ on reducing colonization of EHEC in mice gut intestine. The bacterial numbers of EHEC at 14, 21, and 28 days in ileum (A), jejunum (B), cecum (C), and feces (D) from EHEC-infected mice with no probiotic treatment, LC, or LC^+*mcra*^ pre-treatment were investigated in triplicate. Different letters (‘a’ through ‘c’) at single time point are significantly different (p < 0.05) in the numbers of EHEC among control and treatments.

### Efficient colonization of LC^+*mcra*^ in mice gut

In order to examine the correlation between probiotic colonization and reduction on intestinal bacterial pathogens, we also compared the colonization level of both LC and LC^+*mcra*^ in different portion of mice gut (Fig 4). The one-week daily oral administration led to high and stable cecal colonization level of LC^+*mcra*^ above 10^6^ CFU/g throughout 3 weeks afterwards, which were significantly higher than wild-type LC. A similar trend was found in mice jejunum, whereas, LC^+*mcra*^ only exhibited numerical higher ileum colonization than wild-type LC.

**Fig 4.**
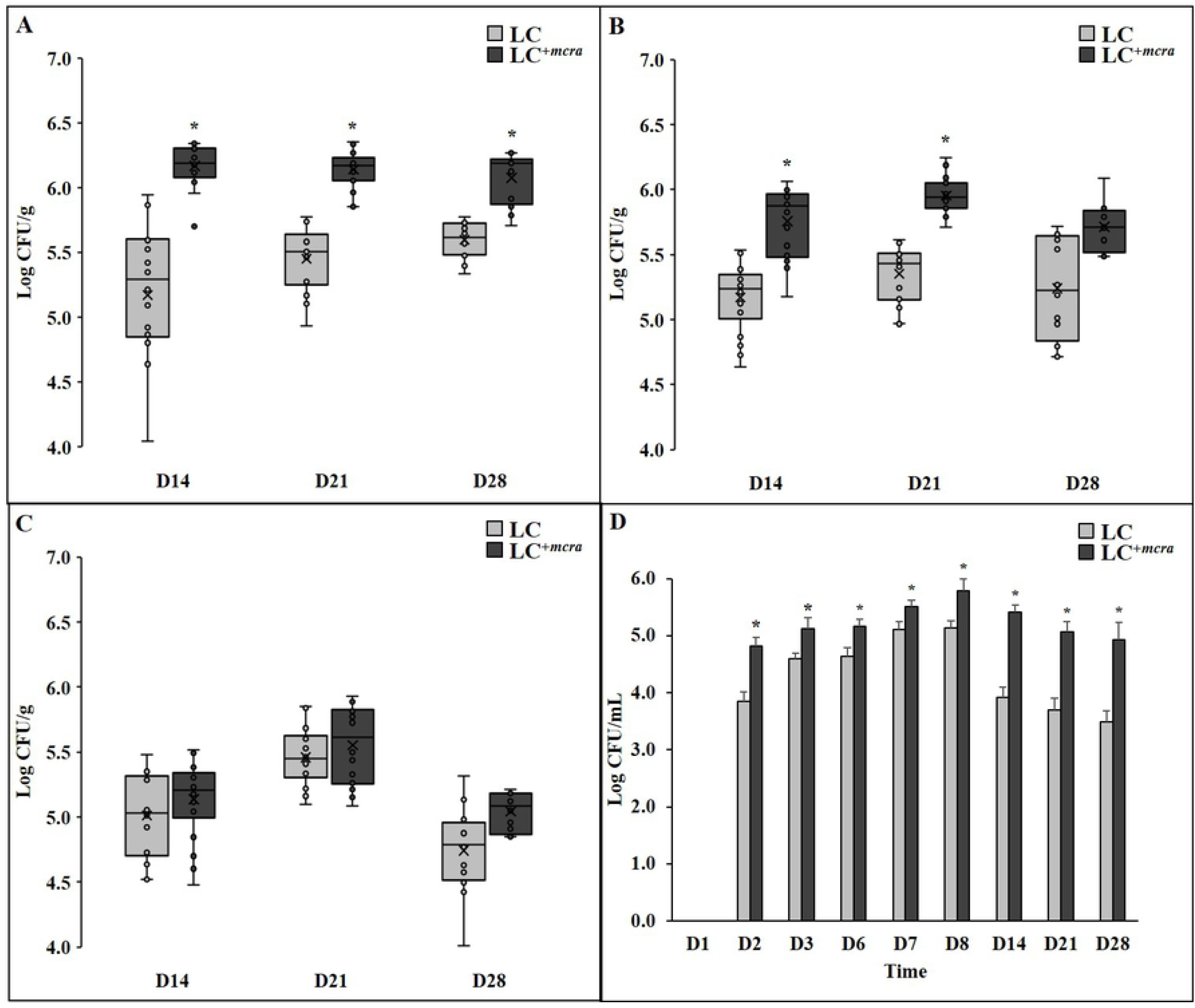
Comparison on colonization levels of LC and LC^+*mcra*^ in mice gut intestine. The bacterial numbers of specific *L. casei* at 14, 21, and 28 days in ileum (A), jejunum (B), cecum (C), and feces (D) from mice daily administered with LC or LC^+*mcra*^ for one week were investigated in triplicate. Asterisk (*) at single time point are significantly different (p < 0.05) in the numbers of gut colonized or fecal shedding wild-type LC and LC^+*mcra*^.

The fecal shedding number of administered LC were observed to raise after 1^st^ day consumption (Fig 4D). Specifically, LC^+*mcra*^ fecal shedding colonies gradually increased from 4.8 log CFU/mL, reached 5.8 log CFU/mL at the next day of final daily administration, and slightly decreased around 5 log CFU/mL after 3 weeks. Whereas, fecal shedding colonies of wild-type LC were observed significantly lower (by 0.4-1.5 log CFU/mL) than LC^+*mcra*^. They reached 5.1 log CFU/mL as peak at the next day of final daily administration and ended up with lower than 3.5 log CFU/mL after 3 weeks.

### Mice hematology

The hematological changes in mice with ST infection with or without pretreated with probiotic strains at various time points were summarized in Table 2. When compared with control group mice with placebo, ST challenge resulted in dramatic increase of red blood cells (RBC) but decrease of white blood cells (WBC) and platelets (PLT). Both pre-treatment of LC or LC^+*mcra*^ alleviated the increment of RBC and loss of WBC/PLT in mice during Salmonellosis. LC^+*mcra*^ treatment on mice could further help the mice maintain their normal levels of RBC, WBC, and PLT.

**Table 1.** Mice groups, numbers per group, and their treatment/infection.

| Group (#) | Mice (n) | Probiotic Treatment<br>(daily during 1<sup>st</sup> week) |  |  | Pathogen challenge<br>(beginning of 2<sup>nd</sup> week) |  |  |
| --- | --- | --- | --- | --- | --- | --- | --- |
|  |  | PBS | LC | LC<sup>+mcrA</sup> | PBS | ST | EHEC |
| A1 | 10 | + | - | - | + | - | - |
| B1 | 10 | - | + | - | + | - | - |
| C1 | 10 | - | - | + | + | - | - |
| A2 | 10 | + | - | - | - | + | - |
| B2 | 10 | - | + | - | - | + | - |
| C2 | 10 | - | - | + | - | + | - |
| A3 | 10 | + | - | - | - | - | + |
| B3 | 10 | - | + | - | - | - | + |
| C3 | 10 | - | - | + | - | - | + |

**Table 2.** Hematological changes and comparison of mice in different groups.

| Day | Red Blood Cells ($10^6/\text{mm}^3$) | | | | White Blood Cells ($10^3/\text{mm}^3$) | | | | Platelets ($10^5/\text{mm}^3$) | | | |
| --- | --- | --- | --- | --- | --- | --- | --- | --- | --- | --- | --- | --- |
|  | Control | Infection | LC | LC<sup>+mcra</sup> | Control | Infection | LC | LC<sup>+mcra</sup> | Control | Infection | LC | LC<sup>+mcra</sup> |
| **14** | 10.15±0.14<sup>*b</sup> | 13.03±4.20<sup>a</sup> | 10.84±0.55<sup>b</sup> | 10.60±0.46<sup>b</sup> | 3.48±0.35<sup>a</sup> | 2.95±0.80<sup>b</sup> | 3.12±0.61<sup>a</sup> | 3.55±0.42<sup>a</sup> | 7±3<sup>a</sup> | 2±2<sup>c</sup> | 5±2<sup>b</sup> | 7±3<sup>a</sup> |
| **21** | 10.34±0.21<sup>b</sup> | 14.89±3.87<sup>a</sup> | 10.72±0.65<sup>b</sup> | 10.51±0.33<sup>b</sup> | 3.65±0.25<sup>a</sup> | 2.96±0.40<sup>b</sup> | 3.13±0.69<sup>a</sup> | 3.54±0.38<sup>a</sup> | 8±1<sup>a</sup> | 2±1<sup>c</sup> | 5±2<sup>b</sup> | 8±3<sup>a</sup> |
| **28** | 10.29±0.13<sup>c</sup> | 15.64±4.37<sup>a</sup> | 11.84±2.14<sup>b</sup> | 10.71±0.53<sup>c</sup> | 3.59±0.22<sup>a</sup> | 2.90±0.49<sup>b</sup> | 3.26±0.30<sup>a</sup> | 3.51±0.29<sup>a</sup> | 7±2<sup>a</sup> | 2±1<sup>c</sup> | 4±2<sup>b</sup> | 8±2<sup>a</sup> |

\* Means with different letters (a-c) for each type of blood cell in different groups at individual time point are significantly different at $p < 0.05$

To further evaluate the WBC composition in blood collected from the mice challenged with ST with or without pre-treated with probiotic strains, we investigated neutrophils, lymphocytes, monocytes, eosinophils, and basophils counts in different time points, which is summarized in Table 3. The numbers of neutrophils and lymphocytes in blood collected from mice challenged with ST were found to be notably reduced, whereas the monocytes, eosinophils, and basophils levels in mice with salmonellosis were detected to be significantly higher. The pre-treatments with either probiotic strain, wild type LC or mutant LC^+*mcra*^ was able to maintain the normal WBC composition under ST infection, including all five cells studied, at the same levels statistically in comparison with control group.

**Table 3.**
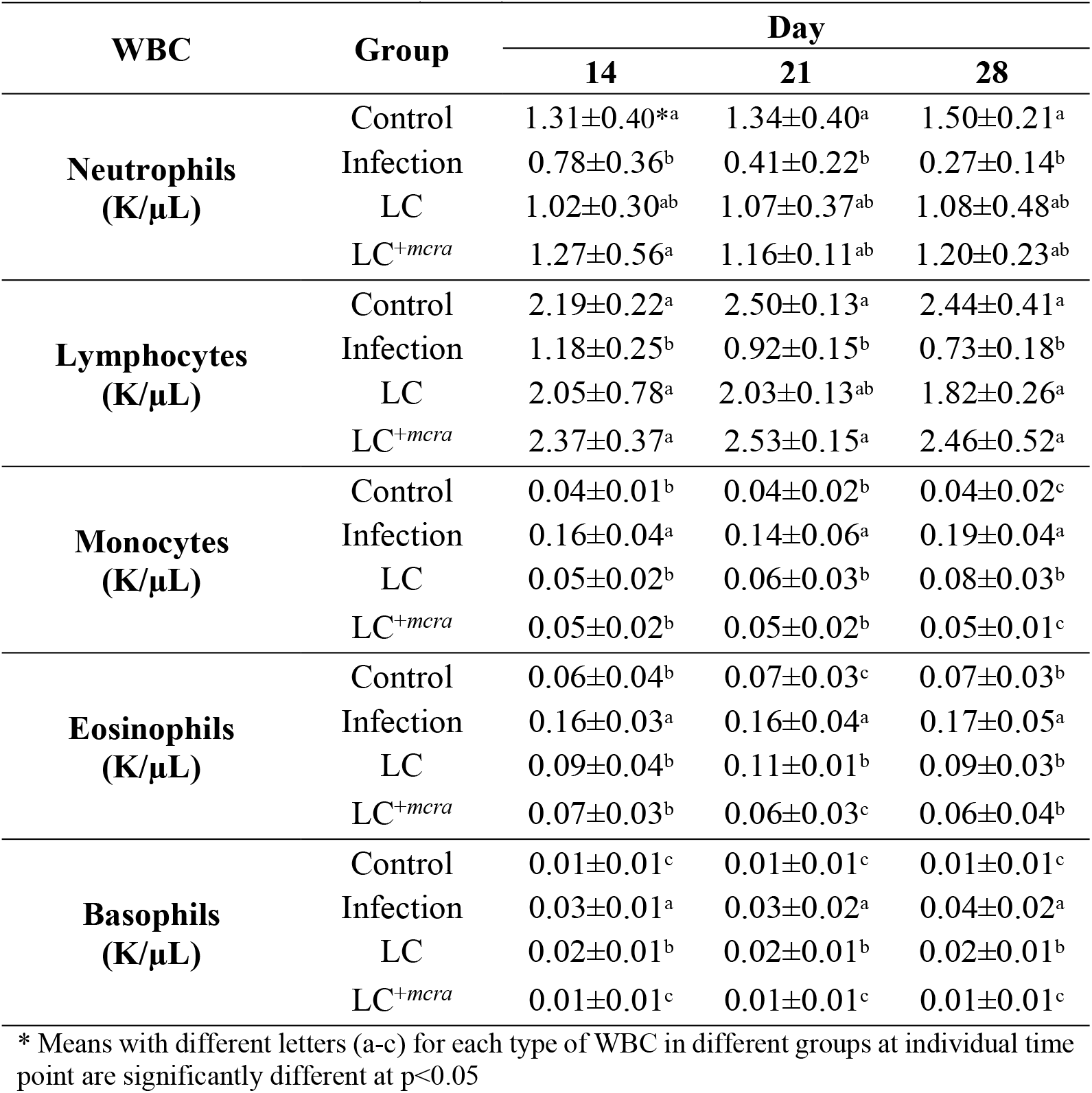
White Blood Cells (WBC) counts of mice in different groups.

### Mice histopathology

The histological examination of mouse cecal sections is shown in Fig 5. Tissue of cecum collected from the control group mice and mice challenged with ST challenge (Fig 5A and 5D) with administration of LC^+*mcra*^ (Fig 5C and 5F) exhibited normal intestinal villi, microvilli, and goblet cells. In comparison, salmonellosis induced variable levels of histological alterations and abnormalities consisting of severe goblet cell depletion, villi/microvilli elimination, and inflammatory infiltrations between circular folds were found in cecum sections from ST infected mice which were not pretreated with probiotics (Fig 5G, 5H, 5I, 5J, 5K, and 5L). However, the tissue of cecum collected from the mice administrated with LC showed symptoms of salmonellosis, but the induced histopathological changes were mild, such as slight goblet cell reduction and slight changes of villi/microvilli (Fig 5B and 5E).

**Fig 5.**
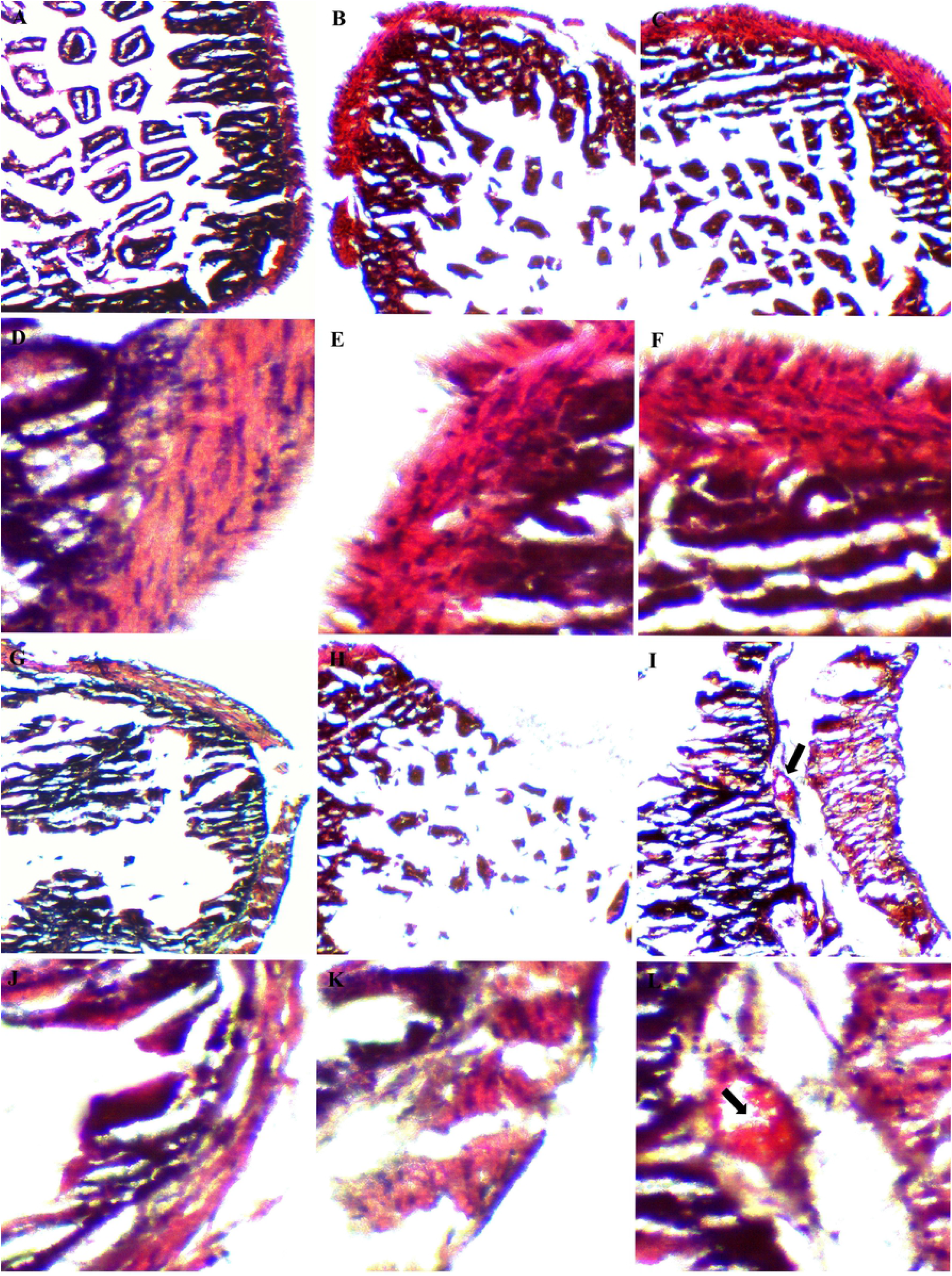
Cecum histopathology in mice. Representative H&E-stained cecum sections from experimental groups were showed in panels (A-C & G-I captured under 100×; D-F & J-L captured under 100×): (A&D) control mice, (B&E) intestinal villi and microvilli reduction in ST-infected mice with 1-week LC pre-treatment, (C&F) normal intestinal histology in ST-infected mice with 1-week LC^+*mcra*^ pre-treatment, (G&J) moderate depletion of goblet cells and villi/microvilli in ST-infected mice, (H&K) massive elimination of goblet cells and villi/microvilli in ST-infected mice, (I&L) intestinal inflammation and infiltration at circular folds in ST-infected mice (arrows).

### Regulation on expression of intestinal inflammatory cytokine genes

The regulation of cecal inflammatory cytokine gene expressions during 3-week ST infection as well as 1-week probiotic pre-administration was displayed in Fig 6. Specifically, ST infection induced up-expression of 4 pro-inflammatory cytokines IL-1β, IL-6, INF-γ, TNF-α genes and 1 anti-inflammatory cytokine IL-10 gene in mice cecal tissue cells. The up-regulation levels ranged from 2.2 to 7.8 log folds with the highest values for INF-γ gene and the highest expression at two weeks after ST challenge (Day 21). Another anti-inflammatory cytokine TGF-β gene was found down-regulated in ST infected mice cecum by 1.5 to 2.9 log folds. The expression of intestinal inflammation-related cytokine genes in LC^+*mcra*^ pre-treated mice were manipulated at a positive manner. For example, all 4 pro-inflammatory cytokine genes provoked by ST were suppressed significantly by 1.3 to 5.3 log folds through three weeks after challenging compared to mice with no probiotic protection; expression of anti-inflammatory cytokines IL-10 and TGF-β genes were stimulated notably in comparison with either control or ST infection with no probiotic prevention.

**Fig 6.**
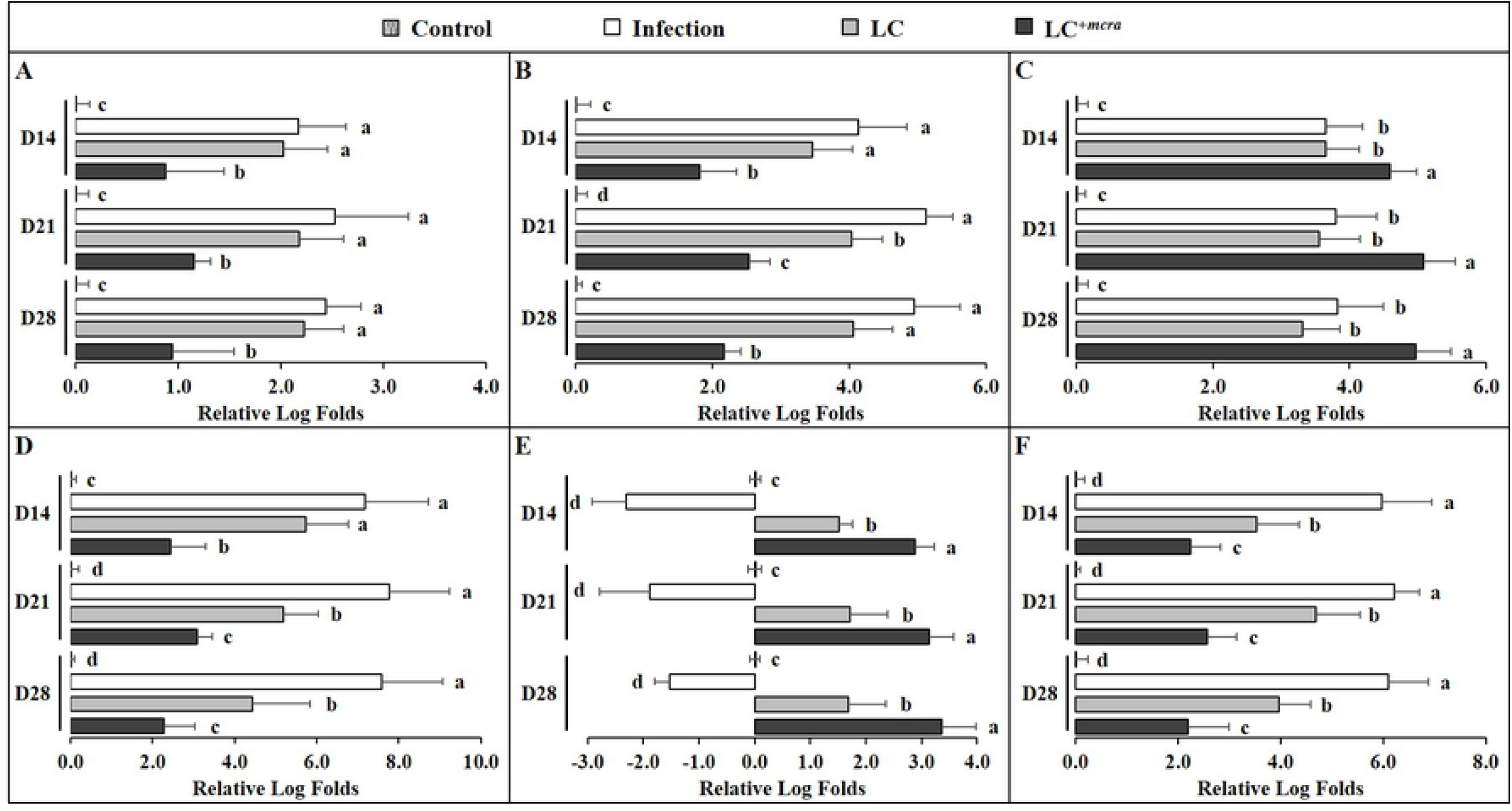
Differential expression levels of mice cecal cytokine genes. The relative log fold changes in expression of IL-1β (A), IL-6 (B), IL-10 (C), INF-γ (D), TGF-β (E), and TNF-α (F) genes from cecum tissue cells collected from mice control, under ST infection, pre-treated with wild-type LC and challenged with ST, or pre-treated with LC^+*mcra*^ and challenged with ST were examined in triplicate. Different letters (‘a’ through ‘d’) at single time point are significantly different (p < 0.05) among control and treatments.

### Modulation on murine gut microbiota composition

To compare the gut microbiome composition in various groups of mice, we randomly selected their cecal contents (5 mice from each group) for 16S metagenomic sequencing and taxonomic classification. According to the taxonomic profile at the phylum level (Fig 7A), Firmicutes were the dominant phylum (63.51%) in mice control group giving placebo (primary control), which was followed by Bacteroidetes (29.37%). The relative abundance of Proteobacteria was 0.92% with individual variation between 0.54% to 1.35%. Significant difference in gut microbial community phylum composition was observed in ST infected mice (Fig 7B), in which group, though the dominant phylum is still Firmicutes (61.26%), the relative abundance of Bacteroidetes was notably decreased to 17.81% and the relative abundance of Proteobacteria boosted to 14.86%. One-week daily administration of probiotic (LC or LC^+*mcra*^) positively shaped the phylum level gut microbiota composition in mice with ST challenging (Fig 7C and 7D). To specify, in comparison with ST infected mice with no probiotic protection (secondary control), the dominance of Firmicutes were raised by 5.67 and 13.34% in LC and LC^+*mcra*^ pretreated groups, respectively. The relative abundances of cecal Proteobacteria were also reduced by 13.17 and 14.17% in LC and LC^+*mcra*^ pretreated mice groups, respectively.

**Fig 7.**
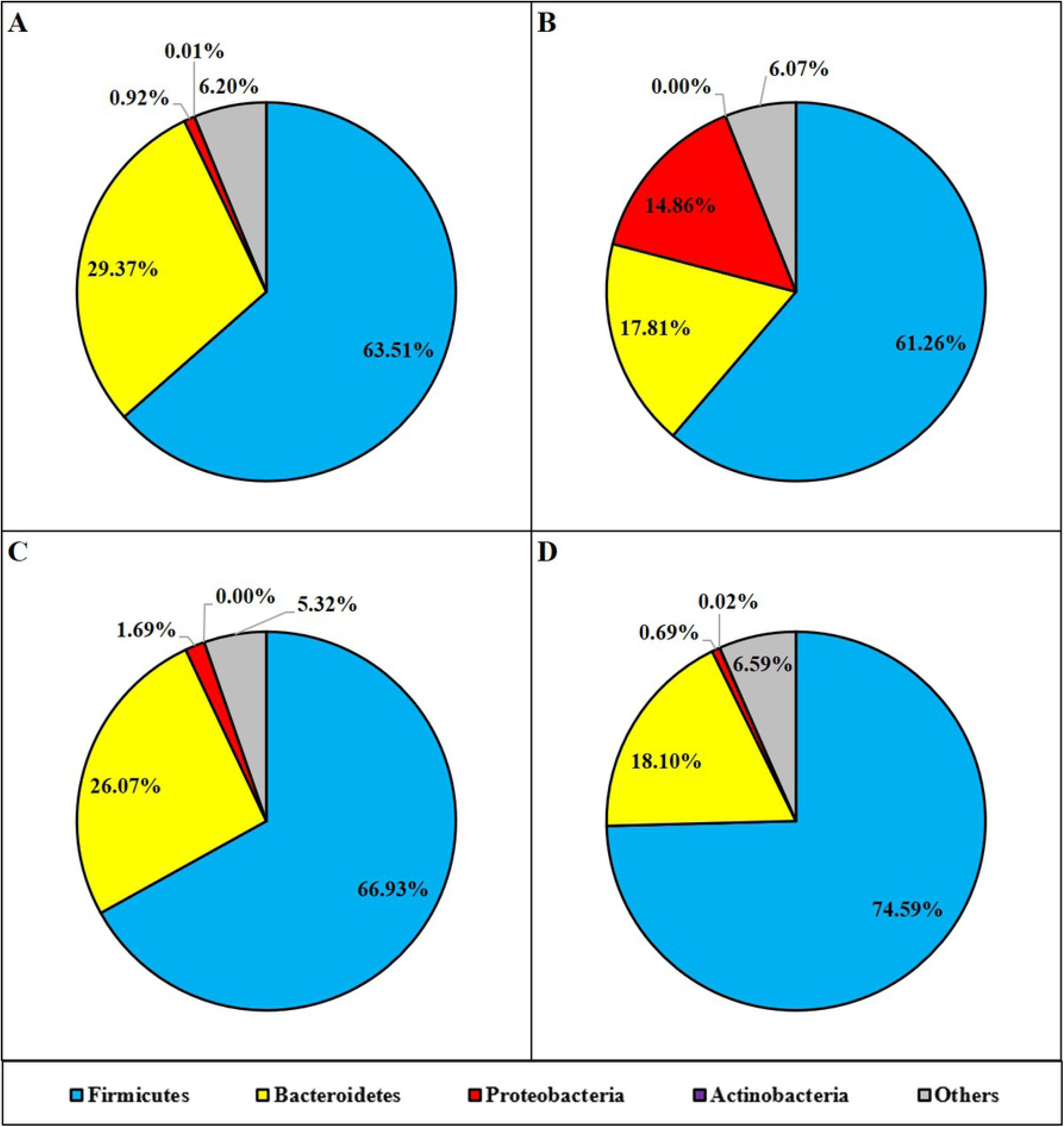
Mice cecal microbial community phylum-level structure. Bacterial distributions at phylum level in cecal contents from individual pooled dataset were depicted in terms of (A) control mice providing placebo, (B) mice infected with ST, (C) mice daily administered with LC for one week followed by ST challenge, and (D) mice daily administered with LC^+*mcra*^ for one week followed by ST challenge.

At genus level (Fig 8), *Bacteroides* was identified being the highest abundant (18.50%) in primary control group of mice cecal contents, followed by *Ruminococcus* (7.17%), *Blautia* (7.02%), *Johnsonella* (4.39%), *Lactobacillus* (1.80%). The relative abundances of *Salmonella* and *Enterobacter* were observed less than 0.01% of the total gut bacterial composition. Whereas, the gut microbiota genus in ST-infected mice exhibited distinctively with significantly higher abundances of *Salmonella* (5.27%) and *Enterobacter* (3.72%), but lower abundances of *Bacteroides* (9.98%), *Blautia* (5.16%), *Johnsonella* (3.34%), and *Lactobacillus* (0.17%) were observed. Other gut microbial genus-level noticeable differences between ST infected mice and control included reduced *Anaerobranca, Anaeroplasma, Butyrivibrio* and raised *Akkermansia, Desulfobacter, Enterococcus, Klebsiella, Leptolyngbya, Natronincola, Staphylococcus,* and *Tolumonas, Trabulsiella*. Compared with secondary control, probiotic pretreatments notably increased the relative abundances of *Bacteroides*, *Blautia*, *Escherichia*, *Johnsonella*, and *Lactobacillus* as well as lowered *Salmonella*, *Enterobacter*, *Klebsiella*, *Tolumonas*, and *Trabulsiella.* Particularly, LC^+*mcra*^ pre-administration in mice modulated the *Salmonella* and *Enterobacter* relative abundances back to control levels in cecum and significantly escalated their relative abundances of *Bifidobacterium* (0.12%), *Blautia* (8.43%), and *Lactobacillus* (9.18%).

**Fig 8.**
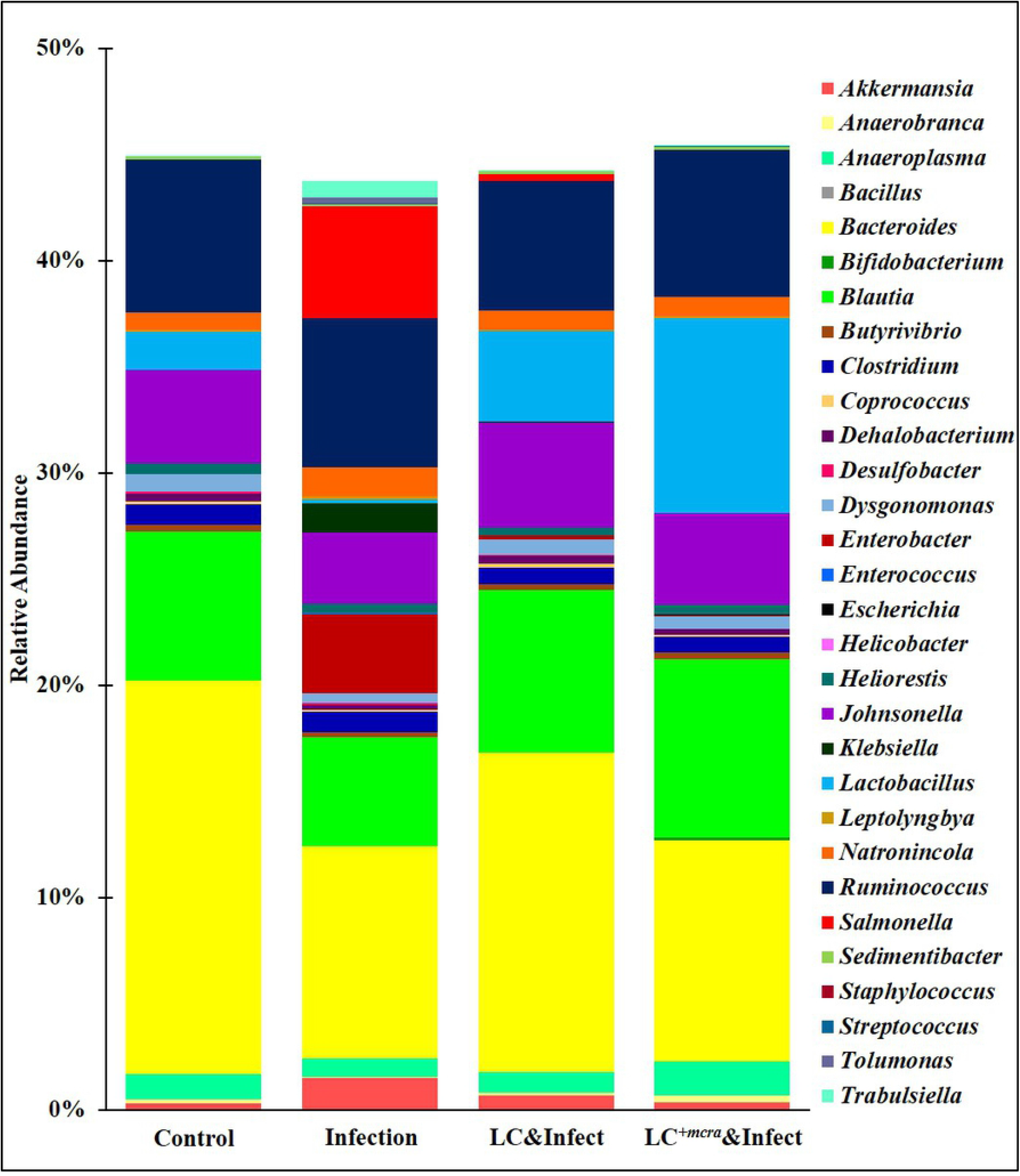
Mice cecal microbiota composition at genus level. Bacterial genus-level community composition in cecal contents from consolidated pool of dataset was compared among different mice groups. Overall 30 bacterial genera were targeted based on their relative abundances and importance in gut microbiome. The total relative abundances of all targeted 30 genera varied from 43 to 46% in different mice groups.

The overall cecal bacterial species diversity was observed to be lowered with ST infection in mice but promoted by LC^+*mcra*^ pre-treatment and protection (Fig 9). Specifically, compared with primary control, the mice group infected with ST exhibited significantly reduced gut intestinal microbial diversity at species level which was indicated by various alpha-diversity indexes including Chao-1, Fisher-alpha, Margalef’s richness, and Simpson (numerically higher), and Shannon. However, the one-week daily pre-administration/prevention with LC^+*mcra*^ instead of with wild-type LC before ST challenging caused a notably increased bacterial species diversity in cecum compared with secondary control group and even higher in comparison with primary control group.

**Fig 9.**
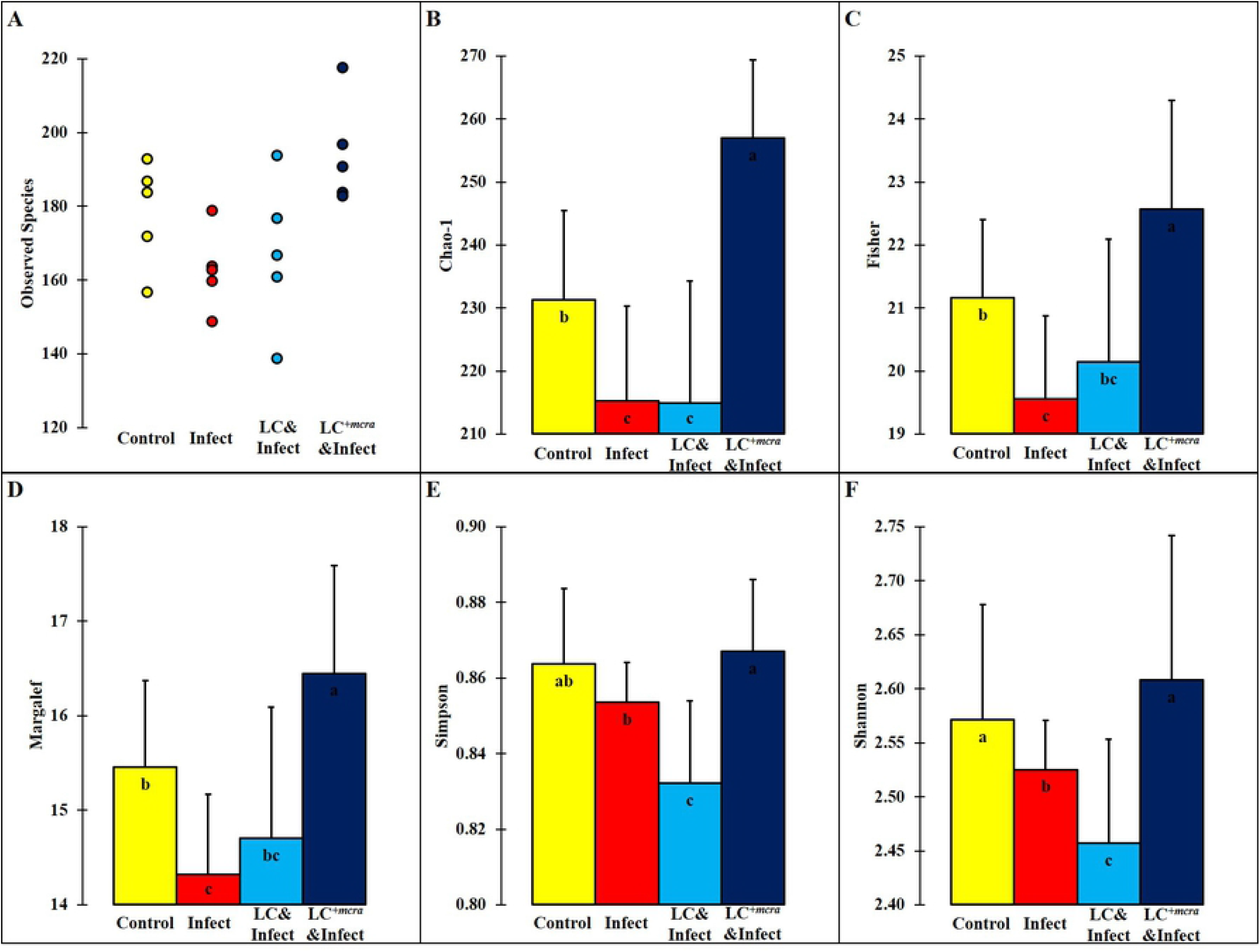
Bacterial diversity at species level in murine cecum. The assessment of alpha-diversity including Observed number of taxa species (A), Chao-1 (B), Fisher’s alpha (C), Margalef‘s richness (D), Simpson index (E), and Shannon index (F) was determined and analyzed among different mice groups. Standard deviations among individual group members were provided. Different letters (‘a’ through ‘c’) are significantly different (p < 0.05) among control and treatments.

## Discussion

The probiotic strain LC^+*mcra*^ with 7-fold upregulation in its expression level of *mcra* gene coding linoleate isomerase has been found with prominently significant 21-fold higher rate in total linoleic acids production per bacterial cell [20]. In a previous study, we revealed *in vitro* that LC^+*mcra*^ could competitively exclude the growth and adhesive activity of both ST and EHEC [20] and at meanwhile, suppress their vital virulence gene factors. Moreover, though effectiveness of probiotics in combatting enteric bacterial pathogens is still controversial, several researchers have suggested that their secondary metabolites such as CLA might enhance their overall *in vivo* health-beneficial functions [14,23–25]. Here in the current study, we systematically and in-depth investigated the double effects of both *Lactobacillus* and CLA on murine gut health. According to our results, 1-week consecutive consumption of LC^+*mcra*^ through oral administration efficiently prevented/mitigated the following *Salmonella* infection. Although probiotic administration through water might generate variance of bio-availability in mice gut, it is worth mentioning that early-staged oral probiotic gavage possesses high risk in potential induced injury in 3-week-old mouse esophagus. The bacterial fecal shedding serves as a key indicator about the gut intestinal colonization [26], correspondingly we observed reduced ST/EHEC in both fecal content and intestinal fluids. Though similar studies conducted based on EHEC were not systematic and completed, *Salmonella* colonization was claimed to be restricted by functional fatty acids oral supplements *in vivo* [25,27–29], in which the virulence gene factors of *Salmonella* were suggested to be manipulated [30,31].

On the other hand, probiotic itself was addressed to be capable of reducing intestinal pathogens through physical repellence and colonization resistance [32–36]. Fortunately, all these studies mentioned above supported our *in vivo* findings in which either wild type or genetically engineered *L. casei* remarkably diminished ST/EHEC colonization in cecum, jejunum, and ileum. Whereas, LC^+*mcra*^ displayed more intensive reductions considering the extraneous strengthening effects implemented by its over-promoted CLA production [14]. In fact, CLA has been documented and linked with antimicrobial active against several enteric bacterial pathogens including *Salmonella* though the specific mechanism are still under study [23,37]. Most importantly, the *in vivo* examination based on BALB/cJ mice model justified the protective roles of LC^+*mcra*^ on combating enteric bacterial pathogens, following and matching with previous *in vitro* outcomes relied on various pathogenic bacterial strains [20,38,39].

In most cases, *Salmonella* infections are associated with diarrhea, weight loss, dramatic alterations in composition of blood cells, as well as death [12,40–42]. Accordantly we detected 10^5^-10^7^ CFU intestinal colonization of ST induced salmonellosis and caused around 8% weight loss, 52% higher level of RBC, 19% and 71% lower levels of WBC (especially neutrophils and lymphocytes) and PLT, and severe cecal inflammation in the survival mice. The physical, hematological, and gut intestinal abnormalities mentioned above in our *in vivo* examination contributed in the 40% death rate of mice challenged with enteric bacterial pathogen ST. However, probiotics in secreting different types of functional fatty acids initiate attenuation in over-reactive gut inflammation through anti-inflammatory activities [14,20,22], which correlates with the LC^+*mcra*^ (CLA) mediated relative up-regulation of murine intestinal anti-inflammatory cytokine genes from mice under salmonellosis found in our study. Therefore, apart from the direct colonization competition and repellence, daily administration of probiotics, especially LC^+*mcra*^, also prevented regular salmonellosis symptoms and maintained the overall physical and gut health condition of mice through mediating immuno-modulation. If in future study, several other tissues including kidney, liver, lung, et al. could be examined for LC^+*mcra*^ pre-treatment on prevention of ST systemic infection.

To address concerns from the host’s point of view, the maintenance of intact and operative gut intestine physiological condition is crucial in both metabolism and symbiotic intestinal microbiota composition [3,43–45]. In our study, LC^+*mcra*^ and its byproduct CLA prevented ST-induced elimination of goblet cells, villi, and microvilli as well as the inflammatory infiltrations between circular folds in cecum, which maintained the overall functions in terms of intestinal nutrients absorption and profoundly raised the survival rate (0 death) in mice. As a matter of fact, CLA has been previously connected with colitis and inflammatory bovine disease recovery [46,47], but the specific mechanisms are still under discovery. Here our findings based on CLA are in support of these researches and suggest a protective mechanism from both bacterial colonization and host histology sides.

A balanced gut microbial ecosystem serves as the crucial defense against colonization and infection with enteric pathogens [14,48,49]. *Salmonella* infection could have negatively impact on gut intestinal microbiome composition by diminishing the abundances of Firmicutes including *Lactobacillus* and *Bifidobacterium,* and simultaneously favoring the dominance of Proteobacteria inducing follow-up opportunistic infections [50–52]. In our study, we observed the raised abundances of *Salmonella* and *Enterobacter* with overall reduced bacterial species diversity following ST challenge in mice, whereas LC^+*mcra*^ pre-administration successfully prevented the negative shifting of gut microbiota composition induced by ST infection. As a matter of fact, CLA-containing diets were reported to alter the fatty acids metabolism and developing homeostatic gut microflora [53,54]. The healthier intestinal microbial distribution shaped by CLA-producing probiotic daily consuming, in terms of higher abundances of *Lactobacillus, Bifidobacterium*, and *Blautia* as well as microbial species diversity/richness, strengthened the first-line gut intestinal defense system against multiple pathogenic bacterial infections, may possess a tight association and be the explanation of reduced bacterial pathogen colonization and inflammation in mice gut.

Based on previous research, EHEC oral challenge on distinct mouse models can result in various levels of colonization, morbidity, and mortality [55]. Specifically, EHEC dose as low as 10^2^ CFU led to cecal colonization and death in germ-free mice [56,57] whereas for conventional mice model like BALB/c, considerably higher dose of EHEC was requisite in order to cause diseases [58,59]. In some cases, infectious dose of EHEC less than 10^10^-10^11^ CFU failed to even introduce cecal colonization [60,61], which parallel with our findings. Based on the current study, 10^7^ CFU EHEC orogastrically challenge on BALB/cJ mice induced 10^2^-10^4^ CFU/g intestinal fluid colonization on cecum, jejunum, and ileum but failed to motivate any visible physiological abnormalities or mortality in mice. This could be explained by the relative resistance in BALB/c mice towards EHEC through shorter shedding duration and producing higher serum/fecal levels of 0157-specific IgA [55,60]. On the other hand, LC^+*mcra*^, as we observed *in vitro* [20] and predicted for *in vivo,* stood out in reducing the colonization level of EHEC as well as preventing from kidney histological abnormalities and weight loss in BALB/cJ mice. Further research dependent on germ-free or compromised commensal flora mouse model might be substantial in revealing how LC^+*mcra*^ involved in defending host from EHEC pathogenesis and post-infectious complications.

To conclude, the current study has demonstrated a substantial influence of CLA over-producing probiotic strain, LC^+*mcra*^ exerted on *Salmonella* and pathogenic *E. coli* infections in conventional mice. Specifically, mice orally given LC^+*mcra*^ daily for one week minimized EHEC colonization and protected themselves from ST-facilitated serious salmonellosis which was observed by notably reduced fecal shedding and intestinal colonization of ST, amelioration on acute inflammation, and prevention on hematological and histological abnormalities. In depth metagenomic analysis revealed that LC^+*mcra*^ pretreatment modulated mice cecal bacterial community with increased diversity which are predominated with comparative higher Firmicutes and lower Proteobacteria. The outstanding protective roles of LC^+*mcra*^ against ST and EHEC infection plus its profound effectiveness over wild-type LC may provide a promising option for prophylaxis on pathogenic *Salmonella* and diarrheagenic *E. coli* infections and reduce enteric bacterial infections.

## Materials and methods

### Ethics statement

Mice *in vivo* experiments were performed in ABSL2 facilities in Department of Animal and Avian Sciences, University of Maryland in accordant with protocol #R-NOV-17-55 approved by the Institutional Animal Care and Use Committee (IACUC). The best effort was made for minimizing the suffer of animals. To ensure animal welfare, mice were monitored and recorded for physical appearance and body weight once/day on a daily basis during experimental period. Animals were euthanized by CO_2_ exposure in a chamber for 5 minutes until all evidences of cardiac function and respiration were absent.

### Bacterial strain and growth conditions

*Lactobacillus casei* (LC, ATCC 334) and our laboratory generated linoleic acid over-expressed *L. casei, LC+^mcra^* [20,38] were used as probiotics while *Salmonella enterica* serovar Typhimurium (ST, ATCC 14028) and enterohemorrhagic *Escherichia coli* EDL933 (EHEC, ATCC700927) were chosen as enteric bacterial pathogens in this study. Both *Lactobacillus* strains were grown on MRS agar at 37 °C for 24 h in the presence of 5% CO_2_ (Forma™ Scientific CO_2_ water jacketed incubator, Thermo Fisher Scientific Inc., Waltham, MA, USA). ST and EHEC were grown on LB agar (EMD Chemicals Inc., Gibbstown, NJ, USA) for 18 h at 37 °C under aerobic conditions (Thermo Scientific MAXQ 4450, Thermo Fisher Scientific Inc., Waltham, MA, USA).

### Mice model and animal experiments

The 3-week-old BALB/cJ Mice (approximately 8-10 g) were purchased from The Jackson Laboratory (Bar Harbor, ME USA) and reared in static micro-isolating cages with cellulose Bio-Performance bedding and huts as environmental enrichment. Teklad standard rodent diet and regular tap water were provided for mice feeding and drinking, respectively. A total of 90 mice (45 male and 45 female) were used for each trial. Following a completely randomized method, 90 mice were randomly assigned to 9 groups (designated A1 to C3) resulting in 10 mice per group; two cages were assigned to each group with a total of 5 mice per cage. Mice cages were changed weekly, and each individual mouse was weighed and monitored with health examinations daily. At the end of the second, third, and fourth week, 3, 3, and 4 mice from each group respectively, were randomly selected and euthanized with CO_2_ inhalation in euthanasia chamber for organ samples collection.

### Feeding probiotic to BALB/cJ mice and challenging with ST and EHEC

Overnight culture of LC or LC^*+mcra*^ in MRS broth were diluted in fresh 5 mL MRS broth at 1:50 and allowed for 3 h further growth. The bacterial cells in exponential phase were harvested following centrifugation at 3,000 × *g* for 15 min, PBS washing, and resuspension in 1.0 mL PBS. A final concentration of 10^11^ CFU/mL was adjusted with PBS and used to feed mice. The design of *in vivo* mouse trial was summarized in Table 1. Probiotic (either 10^9^ CFU/mL LC or LC^+mcra^) cells were maintained in water bottle fill with regular tap water for group B and C and feed to mice from Day 1 to Day 7. Control mice, group A, was fed with regular tap water only.

Overnight culture of ST and EHEC bacterial cells in LB broth were diluted in fresh 5 mL LB broth at 1:50 and allowed for 3-4 h further growth at 37 °C. The exponential phase bacterial cells were harvested and washed by centrifugation at 3,000 × *g* for 15 min and resuspended in 1.0 mL of PBS. A final concentration of bacterial cells was adjusted to 10^8^ CFU/mL in PBS. On Day 7, an aliquot of 100 μL ST or EHEC suspension containing approximately 10^7^ CFU was fed to mice in groups 2 or 3 respectively, with oral gavage, and the mice were reared thereafter for another 3 weeks. Mice in group 1 was orogastrically fed with 100 μL PBS and served as control.

### Sample collection and processing

In order to estimate the bacterial fecal shedding, fecal samples were collected from each mouse in sterile Whirl-Pak bags using sterile spoons at Day 1, 2, 3, 6, 7, 8, 9, 14, 21, and 28 for PBS serial dilution and plating on specific agar plates (MRS agar for *L. casei,* XLT-4 agar for ST, MacConkey agar for EHEC) [62]. In order to investigate the bacterial colonization in mice gut, intestine, ilium, jejunum, and cecum from each euthanized mouse were separated and harvested. Then the ilium, jejunum, and cecal fluids were serial diluted with PBS, followed by plating on specific agar plates. Specifically, MRS agar for *L. casei*, XLT-4 agar for ST, MacConkey agar for EHEC were used, respectively.

Mice cecum was kept in RNA Later for further RNA extraction, cDNA reverse transcription, and inflammation-related gene expression level analysis. For hematological analysis, the blood samples from each mouse was collected from heart in VACUETTER^®^ Heparin tubes (Greiner Bio-One, Monroe, NC, USA) and further analyzed with a ProCyte Dx^®^ Hematology Analyzer (IDEXX, Westbrook, ME, USA) according to the manufacturer’s instructions.

### RNA extraction, cDNA synthesis, and Quantitative RT-PCR for evaluation of targeted gene expressions

Extraction of mice intestinal RNA was carried out using TRIzol^®^ Reagent (Life Technologies Co., Carlsbad, CA, USA) following previous methods [63]. The cDNA synthesis was performed according to the manufacture’s instruction of qScript cDNA SuperMix. The PCR reaction mixture containing 10 μL PerfeCTa SYBR Green Fast Mix (Quanta Biosciences, Beverly, MA, USA), 2 μL of each 100 nM primer, 2 μL of cDNA (10 ng), and 4 μL of RNase-free water was amplified using an Eco Real-Time PCR system with 30 sec denaturation at 95 °C, followed by 40 cycles of 95 °C for 5 sec, 55 °C for 15 sec, and 72 °C for 10 sec. All the relative transcription levels of target genes were estimated by comparative fold change. The CT values of genes were normalized to the housekeeping gene, and the relative expression levels of target genes were calculated by the comparative method [64]. Quantitative RT-PCR was carried out in triplicate.

### Histopathology analysis

Intestinal tissue samples were taken from mice after euthanization and were stored in neutral buffered formalin (4% formaldehyde; pH 7.4) at 4°C for further processing. Once the samples were removed from fixative, they were dehydrated with increasing concentrations of ethanol, cleared in xylene, and embedded in paraffin. Microtome (LEICA RM2065, Leica Biosystems, Buffalo Grove, IL, USA) was used to harvest 5 μm thick paraffin sections followed by heat fixing at 37 °C overnight. Then the slices were stained with hematoxylin and eosin and mounted with DPX mounting medium 13512 (Electron Microscopy Sciences, Hatfield, PA, USA). Histological observations were performed under a light microscope (BA210E, Motic Asia, Hong Kong, China).

### Metagenomic sequencing and analysis

Mice cecal contents were harvested and 5 samples from each group of control, ST infection, LC pretreatment followed by ST infection, or LC^+*mcra*^ pretreatment followed by ST infection were randomly selected for metagenomics analysis. Microbial genomic DNA extraction was carried out using QIAamp Fast DNA Stool Kit (QIAGEN, Valencia, CA, USA) following instructions from the manufacturer. The variable V3 and V4 regions of microbial 16S rRNA gene were targeted for phylogenetic classifications. DNA libraries were prepared for equimolar-pooling using Nextera DNA Library Preparation Kit and Nextera Index Kit (Illumina, San Diego, CA, USA) according to the manufacturer’s instructions. Paired-end sequencing (2 × 300 bp) was conducted on Illumina MiSeq using MiSeq v3 600-cycle kit (Illumina, San Diego, CA, USA). Sequence data was processed through MiSeq Reporter - BaseSpace for FASTQ Workflow generation followed by taxonomic classification based on Greengenes database (http://greengenes.lbl.gov/). Demultiplexing was performed using only perfect index recognition (mismatch = 0) followed by removing PhiX reads. 16S sequence length below 1250 bp or with more than 50 wobble bases was filtered, and all entries classified with no genus or species were also filtered. The relative abundances and alpha-diversity indices were calculated using ‘vegan’ R package and plotted in Excel.

### Statistical analysis

All data were analyzed by the SPSS software. Comparison among multiple mice groups were performed with the one-way analysis of variance followed by Tukey’s and Bonferroni’s tests. For all tests, significant differences were considered on the basis of P values below a significant level of 0.05.

## Acknowledgements

The authors would like to thank Dr. Vinod Nagarajan, Zabdiel Alvarado Martinez, Arpita Aditya, for their assistance during animal experiments. We also thank Dr. Rachel Dennis and Jasmine Mengers from department of Animal and Avian Sciences in the University of Maryland for their guide on histopathological studies.

## Author Contributions

**Conceptualization**: Mengfei Peng, Debabrata Biswas.

**Data curation**: Mengfei Peng, Zajeba Tabashsum.

**Formal analysis**: Mengfei Peng.

**Investigation**: Mengfei Peng, Zajeba Tabashsum, Debabrata Biswas.

**Methodology**: Mengfei Peng, Zajeba Tabashsum, Puja Patel, Cassandra Bernhardt.

**Resources**: Jianghong Meng, Debabrata Biswas.

**Supervision**: Jianghong Meng, Debabrata Biswas.

**Writing – original draft**: Mengfei Peng.

**Writing – review & editing**: Mengfei Peng, Chitrine Biswas, Debabrata Biswas.

